# Cryopreservation of neuroectoderm on a pillar plate and *in situ* differentiation into human brain organoids

**DOI:** 10.1101/2024.07.25.605147

**Authors:** Mona Zolfaghar, Prabha Acharya, Pranav Joshi, Na Young Choi, Sunil Shrestha, Vinod Kumar Reddy Lekkala, Soo-Yeon Kang, Minseong Lee, Moo-Yeal Lee

## Abstract

Cryopreservation in cryovials extends cell storage at low temperatures, and advances in organoid cryopreservation improve reproducibility and reduce generation time. However, cryopreserving human organoids presents challenges due to the limited diffusion of cryoprotective agents (CPAs) into the organoid core and the potential toxicity of these agents. To overcome these obstacles, we developed a cryopreservation technique using a pillar plate platform. To illustrate cryopreservation application to human brain organoids (HBOs), early-stage HBOs were produced by differentiating induced pluripotent stem cells (iPSCs) into neuroectoderm (NEs) in an ultralow atachement (ULA) 384-well plate. These NEs were transferred and encapsulated in Matrigel on the pillar plate. The early-stage HBOs on the pillar plate were exposed to four commercially available CPAs, including PSC cryopreservation kit, CryoStor CS10, 3dGRO, and 10% DMSO, before being frozen overnight at -80°C and subsequently stored in a liquid nitrogen dewar. We examined the impact of CPA type, organoid size, and CPA exposure duration on cell viability post-thaw. Additionally, the differentiation of early-stage HBOs on the pillar plate was assessed using RT-qPCR and immunofluorescence staining. The PSC cryopreservation kit proved to be the least toxic for preserving these HBOs on the pillar plate. Notably, smaller HBOs showed higher cell viability post-cryopreservation than larger ones. An incubation period of 80 minutes with the PSC kit was essential to ensure optimal CPA diffusion into HBOs with a diameter of 400 - 600 µm. These cryopreserved early-stage HBOs successfully matured over 30 days, exhibiting gene expression patterns akin to non-cryopreserved HBOs. The cryopreserved early-stage HBOs on the pillar plate maintained high viability after thawing and successfully differentiated into mature HBOs. This on-chip cryopreservation method could extend to other small organoids, by integrating cryopreservation, thawing, culturing, staining, rinsing, and imaging processes within a single system, thereby preserving the 3D structure of the organoids.

## Introduction

The development of 3D cell/tissue models has revolutionized drug screening, disease modeling, and personalized medicine (1). Despite the growing scientific interest in 3D cell models such as organoids, comprehensive investigation of cryopreservation methods for human organoids remain underexplored. Cryopreservation is crucial for the long-term storage of organoids and significantly advances research in the field of 3D cell/tissue models (2). Given the complexity and the long-term culture required for organoid generation, effective cryopreservation could alleviate the need for continuous culture of organoids, conserving resources and enhancing organoid biobanking (3).

Human brain organoids (HBOs) effectively replicate many aspects of fetal human brains, such as cell diversity, cytoarchitecture, and cellular interactions (4). The cryopreservation of HBOs is highly valuable in the fields of organoid research and neuroscience, particularly for scientists who need to conduct repeated experiments on the same tissue samples over long durations. There are two primary methods of organoid cryopreservation: vitrification and slow freezing. Slow freezing, the traditional approach, uses lower cryoprotectant agent (CPA) concentrations (5 - 10%) and a gradual cooling rate^5^. In contrast, vitrification involves higher CPA concentrations (30 - 50%) and a rapid cooling rate ^5,6^. CPAs are crucial in both processes, as they inhibit ice crystal formation and facilitate thorough penetration into the organoids ^7^.

Nonetheless, both slow freezing and vitrification methods present limitations for CPA penetration. Slow freezing limits CPA penetration due to its lower concentration, whereas the higher concentration in vitrification can be toxic and increase osmotic stress ^8–10^. Achieving the correct CPA balance and choosing a suitable cryopreservation technique are crucial for preserving the structural and functional integrity of organoids during freeze-thaw cycles. Cryoinjury, caused by ice crystal formation, osmotic stress, and temperature fluctuations, poses a significant challenge. CPAs act as a substitute for water in cells, preventing ice crystal formation and maintaining osmotic balance during freezing. However, the diffusion of CPAs in 3D cell/tissue models is limited by the structural and physiological characteristics of these models ^11^. Optimal cryopreservation of organoids necessitates careful consideration of various factors, including CPA selection, incubation time, and temperatures before and after freezing ^8,12^. Ensuring comprehensive CPA diffusion is critical to maintain organoid viability throughout these processes.

In this study, we focus on optimizing the diffusion of CPAs in two different organoid sizes. Additionally, to minimize mechanical damage and stress, and to enhance cryopreservation, we introduce an *in-situ* slow freezing method for early-stage, human brain organoids (ES-HBOs) using a 36-pillar plate with sidewalls and slits (36PillarPlate). The pillar plate, a microfabricated culture device previously employed in our 3D cell culture studies, features 36 vertical pillars, each supporting an individual organoid ^13–15^.

Cryopreserving with pillar plates eliminates the need for harvesting and centrifugation, allowing for easy CPA removal through gentle washing. The design ensures uniform and swift warming of organoids, reducing ice crystal formation and cell injury. In addition, the architecture prevents organoid aggregation, facilitating their imaging and monitoring in an organized manner. Successful cryopreservation maintains cell and organoid viability, proliferation, and function ^16^. Freeze-thawed ES-HBOs on the pillar plate retained their morphology, exhibited high viability, and eventually differentiated into mature brain organoids.

## Materials and Methods

### Culture of human induced pluripotent stem cells (iPSCs)

Human induced pluripotent stem cell (iPSC) lines were acquired from the Cedar Sinai Biomanufacturing Center and cultured in 6-well plates pre-coated with 0.5 mg of hESC-qualified Matrigel (Corning, Cat. No. 356234). These cells were maintained in a 5% CO_2_ incubator at 37°C and were fed every 24 to 48 hours with mTeSR Plus medium (StemCell Technologies, 100-0276). The iPSCs were passaged using the StemPro EZPassage tool (Life Technologies, 23181-010) upon reaching 70 to 80% confluency, which exhibited less than 10% differentiation. Two iPSC lines, EDi029A and EDi027A, were used in this study, derived from male and female tissues, respectively.

### Formation of embryoid bodies (EBs)

Colonies of human iPSCs were dissociated into single cells using Accutase (Gibco, A1110501) at 37°C for 8 - 10 minutes. After dissociation, single cells were resuspended by gentle pipetting and transferred to a sterile conical tube. The cells were then centrifuged at 300 x g for 5 minutes, and the supernatant was discarded. The cell pellet was mixed with 1 mL of embryoid body (EB) formation medium prepared according to the Lancaster protocol ^17^. The EB formation medium comprised of 40 mL of DMEM-F12 (ThermoFisher, 11320033), 10 mL of knockout serum replacement (KOSR; ThermoFisher, 10828010), 1.5 mL of ESC-quality FBS (ThermoFisher, 10439001), 500 μL of GlutaMAX (Invitrogen, 35050-038), 500 μL of MEM-nonessential amino acids (MEM-NEAA; MilliporeSigma, M7145), 4 ng/mL basic fibroblast growth factor (bFGF; Peprotech, 100-18B), and Rho kinase (ROCK) inhibitor. In this study, the ROCK inhibitor was replaced with a CEPT cocktail consisting of 5 µM Emricasan (Fisher Scientific, 50-136-5235), 0.7 µM Trans-ISRIB (Tocris, 5284), 50 nM Chroman 1 (Tocris, 7163), and a polyamine supplement diluted at 1:1,000 (MilliporeSigma, P8483). The cell counting was conducted using Trypan blue and an automated cell counter (BIO-RAD, TC20). Cells suspended in the EB formation medium were seeded into an ultralow attachment (ULA) 384-well plate (S-bio, 20939308) at seeding densities of 1,000 and 3,000 cells per well, each containing 50 µL of the EB formation medium. After 24 hours of culture, small spheroids with a distinct outer layer were observed. The cells were cultured in the EB formation medium supplemented with CEPT and bFGF until day 4, after which the medium was replaced with the EB formation medium without CEPT and bFGF for one additional day. The culture medium was refreshed every other day by removing half of the spent medium and replenishing it with an equal volume of fresh medium. The EB formation process was completed within 5 days of culture in the EB formation medium.

### Neuroectoderm (NE) formation

To generate neuroectoderm (NE), EBs were cultured in neural induction medium (NIM) for two additional days within the same ULA 384-well plate. This was achieved by carefully aspirating most of the EB formation medium and then adding the NIM. The NIM is composed of DMEM-F12, 1% (vol/vol) N2 supplement (ThermoFisher, 17502048), 1% (vol/vol) GlutaMAX, 1% (vol/vol) MEM-NEAA, and 1 µg/mL heparin (MilliporeSigma, H3149) (18). NEs were cultured until day 7, after which they were transferred to a 36PillarPlate preloaded with 5 µL of Matrigel.

### NE transfer from the ULA 384-well plate to the 36PillarPlate preloaded with Matrigel

On day 7, NEs in the ULA 384-well plate were transferred to the 36PillarPlate (Bioprinting Laboratories Inc., 36-01-00) by a sandwiching and inverting method ^15^. The 36PillarPlate, manufaced by injection molding of polystyrene, feastures a 6 x 6 array of pillars with a 4.5 mm pillar-to-pillar distance, 11.6 mm pillar height, and 2.5 mm outer and 1.5 mm inner diameter of pillars ^13,14,18^. Each pillar is uniquely designed with sidewalls and slits suitable for culturing a single NE or organoid encapsulated in Matrigel. For the transfer and encapsulation process, the pillar plate with 5 µL of 6 - 8 mg/mL Matrigel was sandwiched onto the ULA 384-well plate containing NEs. This assembly was then inverted and incubated for 30 - 40 minutes at 37°C in a CO_2_ incubator. Subsequently, the 36PillarPlate with NEs was detached from the ULA 384-well plate and combined with a clear-bottom 384-deep well plate (384DeepWellPlate), also manufactured by injection molding of polystyrene (Bioprinting Laboratories Inc., 384-02-00), which was filled with 80 µL of the neural induction medium (NIM) per well (**Figure 1**). The 384DeepWellPlate is designed with a 16 x 24 array of deep wells, measuring 3.5 mm in width, 3.5 mm in length, and 14.7 mm in depth, with a 4.5 mm distance between wells, to facilitate static cell culture.

### Assessing basal cytotoxicity of CPAs

The NEs on the 36PillarPlate were cultured in NIM for an additional day. On day 8, NEs on the pillar plate were exposed to three commercially available CPAs, including 3dGRO (Sigma-Aldrich, SCM301), PSC cryopreservation kit (Gibco, A2644601), and CryoStor CS10 (StemCell Technologies, 100-1061), as well as a custom-made freezing solution composed of 10% dimethyl sulfoxide (DMSO; Sigma-Aldrich, D8418) in DMEM/F12. The 36PillarPlate containing NEs was then sandwiched onto a 384DeepWellPlate with 50 µL/well of the respective CPAs, and this assembly was incubated at room temperature for 2 hours. Subsequently, the basal cytotoxicity of the CPAs against the NEs was assessed by rinsing the NEs with phosphate-buffered saline (PBS, Gibco, 10010031) twice, followed by incubation with the CellTiter-Glo 3D cell viability assay kit (Promega, G968B) (**Figure 2**). NEs on the pillar plate without CPA exposure served as a control.

### Molecule diffusion into the core of NEs

To indirectly estimate the rate of CPA diffusion into the NE core, NEs on the 36PillarPlate were incubated with CPA and 1 µM calcein AM (ThermoFisher, L3224) at 50 µL/well in a 384DeepWellPlate at room temperature for up to 2 hours. Changes in green fluorescence were monitored using a fluorescence microscope (Keyence, BZ-X810) to determine the diffusion rate of calcein AM into the NE core (**Figure 3**). It was assumed that the diffusion rates of CPA and calcein AM are similar, as both are small molecules.

For the optimal CPA selected (the PSC cryopreservation kit), NEs on the pillar plate were incubated with the CPA for various time intervals: 0, 20, 40, 60, 80, 100, and 120 minutes. After being rinsed with PBS twice, the NEs were further incubated for 24 hours in NIM to accurately measure cell viability. The viability of the CPA-exposed NEs was determined using the CellTiter-Glo 3D cell viability assay kit (**Figure 4**). NEs on the pillar plate without CPA exposure served as a control.

### Determining optimal cell seeding density for NE cryopreservation

To test the optimal cell seeding density for cryopreservation, day 8 NEs on the 36PillarPlate, derived from initial seeding densities of 1,000 and 3,000 cells, were incubated with the PSC cryopreservation kit in the 384DeepWellPlate for 90 minutes at room temperature. Following this, the pillar plate containing day 8 NEs was separated and sandwiched onto an empty 384DeepWellPlate. As the NEs in Matrigel on the pillar plate absorbed the CPA, no excess CPA was required for cryopreservation. For slow freezing, the sandwiched plates with CPA-treated NEs were placed in a Mr. Frosty freezing container (ThermoFisher, 5100-0001) and stored at - 80°C overnight. The plates were then transferred to a liquid nitrogen dewar for 3 days. After storage in liquid nitrogen, the pillar plate with frozen NEs was thawed at room temperature for 2 minutes. To eliminate any residual PSC cryopreservation kit frp, the NEs, the pillar plate was rinsed with pre-warmed NIM containing CEPT in a 384DeepWellPlate, three times for 10, 25, and 45 minutes, respectively. The cryopreserved and thawed NEs on the pillar plate were subsequently cultured in NIM supplemented with RevitaCell™ (a component of the PSC cryopreservation kit, ThermoFisher) for 24 hours. Finally, the viability of the NEs was assessed using the CellTiter-Glo 3D cell viability assay kit (**Figure 5**).

### Cryopreservation of early-stage human brain organoids (ES-HBOs)

The optimal protocols for the formation of ES-HBOs on the pillar plate, on-chip cryopreservation using the PSC cryopreservation kit, and the maturation of cryopreserved ES-HBOs are detailed in **Figure 6**. To generate uniform cerebral organoids, we used the STEMdiff™ Cerebral Organoid Kit (Cat No. 08570), which is based on the formulation published by Lancaster et al ^17^. In summary, the iPSCs were seeded in a ULA 384-well plate at a seeding density of 1,000 cells/well and cultured for 5 days in STEMdiff organoid formation medium, supplemented with the CEPT cocktail for the initial 4 days. After formation, the EBs in the ULA 384-well plate were cultured for an additional 2 days in STEMdiff neural induction medium (NIM). The NEs formed and on day 7 were then transferred to a 36PillarPlate with 5 µL of 6 - 8 mg/mL Matrigel per pillar using the sandwiching and inverting method. Finally, the 36PillarPlate containing NEs was incubated for one more day in STEMdiff NIM.

On day 8, the pillar plate containing NEs was sandwiched onto a 384DeepWellPlate with the PSC cryopreservation kit and incubated for 40 minutes at room temperature. Subsequently, the pillar plate with CPA-exposed NEs was combined with an empty 384DeepWellPlate, sealed, and stored in a Mr. Frosty freezing container at - 80°C overnight. It was then transferred to a liquid nitrogen dewar for 3 days. A minimum of 60 minutes of incubation with the PSC cryopreservation kit is required for complete diffusion of the CPA into the NE core at this cell density. Therefore, the remaining incubation time was completed in the Mr. Frosty freezing container, cooling at a rate of -1°C/min.

After the cryopreservation process in the liquid nitrogen dewar, the pillar plate containing the frozen NEs was thawed at room temperature for 2 minutes. To eliminate any remaining PSC cryopreservation kit residue, the pillar plate underwent three consecutive rinses of 10, 25, and 45 minutes with pre-warmed NIM supplemented with the CEPT cocktail, using a 384DeepWellPlate. The cryopreserved and thawed NEs on the pillar plate were then cultured in STEMdiff neural expansion medium with RevitaCell™ supplement for 1 day. The viability of the NEs was evaluated using the CellTiter-Glo 3D cell viability assay kit, along with calcein AM and ethidium homodimer-1 (EthD-1) staining at 1 and 8 days post-thaw. Additionally, the NEs on the pillar plate underwent further differentiation in NEM with the CEPT cocktail for 5 days. In the final step, the NEs were differentiated for 25 days in STEMdiff maturation medium to form brain organoids.

### Analysis of neural gene expression in brain organoids by RT-qPCR

To investigate the impact of cryopreservation on organoid differentiation and maturation, we compared the expression levels of neural genes in day 38 brain organoids derived from cryopreserved NEs with those from non-cryopreserved brain organoids and induced pluripotent stem cells (iPSCs) as controls using reverse transcription quantitative polymerase chain reaction (RT-qPCR). Brain organoids embedded in Matrigel on the 36PillarPlate were isolated using the Cultrex organoid harvesting solution (R&D Systems, 3700-100-01), which facilitates the non-enzymatic depolymerization of Matrigel. The 36PillarPlate containing brain organoids was sandwiched with a 384DeepWellPlate filled with 80 µL of Cultrex organoid harvesting solution. The sandwiched plates were incubated for 30 minutes at 4°C and then centrifuged at 100 RCF for 10 minutes to separate the organoids. Total RNA was extracted from both iPSCs and brain organoids using the RNeasy Plus Mini Kit (Qiagen, 74134) following the manufacturer’s protocol. cDNA was synthesized from 1 µg of RNA using the protocol provided with the cDNA conversion kit (Applied Biosystems, 4368814). Real-time PCR was performed with the SYBR™ Green Master Mix (ThermoFisher, A25742) and specific forward/reverse primers from IDT Technology (**Supplementary Table 1**). The PCR cycle included 40 repetitions at 95°C for denaturation (30 seconds), annealing at 58 - 62°C (45 seconds, depending on the primer pair), and extension at 72°C (30 seconds) using the QuantStudio™ 5 Real-Time PCR System (Applied Biosystems, A28574). Expression levels of the targeted genes were normalized to the housekeeping gene, glyceraldehyde 3-phosphate dehydrogenase (GAPDH). Analyzed neural genes included *CTIP2* (Coup-TF Interacting Protein 2), *FOXG1* (Forkhead Box G1), *MAP2* (Microtubule-Associated Protein 2), *PAX6* (Paired Box 6), *SOX2* (SRY-Box Transcription Factor 2), *TBR2* (T-box Brain Protein 2), *TUBB3* (Tubulin Beta 3 Class III), and the pluripotency marker *OCT4* (Octamer-Binding Transcription Factor 4).

### Immunofluorescence (IF) staining of brain organoids on the pillar plate

For *in situ* immunofluorescence (IF) staining of brain organoids on the pillar plate, all solutions and reagents used for staining were dispensed at 80 µL/well into the 384DeepWellPlate. Brain organoids on the pillar plate were rinsed three times with PBS for 10 minutes each and then fixed with 4% paraformaldehyde (PFA, ThermoFisher, J19943.K2) in the deep well plate overnight at 4°C. After rinsing three times with PBS for 10 minutes each, the organoids were permeabilized with 0.5% Triton X (Sigma, 9036195) in PBS for 1 hour at room temperature and then exposed to a blocking buffer (consisting of 1% bovine serum albumin (BSA) in PBS with 0.5% Triton X) overnight at 4°C. Primary antibodies were diluted to recommended concentrations using the blocking buffer and the organoids were incubated with them overnight at 4°C. Following primary antibody staining, the organoids were rinsed three times with 0.2% Triton X in PBS for 15 minutes each and incubated with appropriate secondary antibodies diluted in the blocking buffer for 2 - 4 hours at room temperature on a rocker. The stained organoids were then rinsed with 0.5% Triton X in PBS three times for 15 minutes each on a rocker and incubated with 1 µg/mL DAPI in 0.5% Triton X in PBS for 45 minutes. This was followed by a final wash with PBS three times for 10 minutes each and incubation with a tissue clearing solution (Visikol Histo-M, HH-10) for 1 hour. Images were captured with a confocal microscope (LSM710, Zeiss) at 10x and 40x magnification and analyzed with ImageJ/Fiji v1.54f software. The primary and secondary antibodies used are detailed in **Supplementary Tables 2 and 3**, respectively.

### Statistical analysis

Statistical analysis was performed using GraphPad Prism 9.3.1 (GraphPad Software). All experiments were carried out in triplicate and the data are presented as mean ± SD. P values were calculated using t-tests and one-way ANOVA. The statistical significance threshold was set at **** for p < 0.0001, *** for p < 0.001, ** for p < 0.01, * for p < 0.05, and ns = not significant (p > 0.05). Sample sizes are indicated in the figure legends.

## Results

### Basal toxicity of cryoprotective agents (CPAs) against neuroectoderm (NE)

On day 7, NEs formed in a ULA 384-well plate at seeding densities of 1,000 and 3,000 cells/well were transferred to a 36PillarPlate with Matrigel using the sandwiching and inverting method (**Figure 1**). To select the optimal CPA with minimal cytotoxicity, the basal toxicity of the CPAs against NEs was assessed by measuring cell viability after CPA exposure for 2 hours using the CellTiter-Glo 3D assay kit (**Figure 2**). The pillar plate with NEs was sandwiched onto the 384DeepWellPlate containing the PSC cryopreservation kit, CryoStor CS10, 3dGRO, and 10% DMSO at room temperature for 2 hours. The viability of NEs exposed to the CPAs was compared to a control group of NEs not treated with any CPAs. For NEs prepared with an initial cell density of 1,000 cells, the average viability when treated with the PSC cryopreservation kit, 10% DMSO, 3dGRO, and CryoStor CS10 was 90%, 80%, 60%, and 43%, respectively. Similarly, for NEs with an initial cell density of 3,000 cells, the average viability with the PSC cryopreservation kit, 10% DMSO, 3dGRO, and CryoStor CS10 was 98%, 100%, 60%, and 50%, respectively (**Figure 2**). Therefore, the PSC cryopreservation kit exhibited minimal toxicity against NEs.

**Figure 1.**
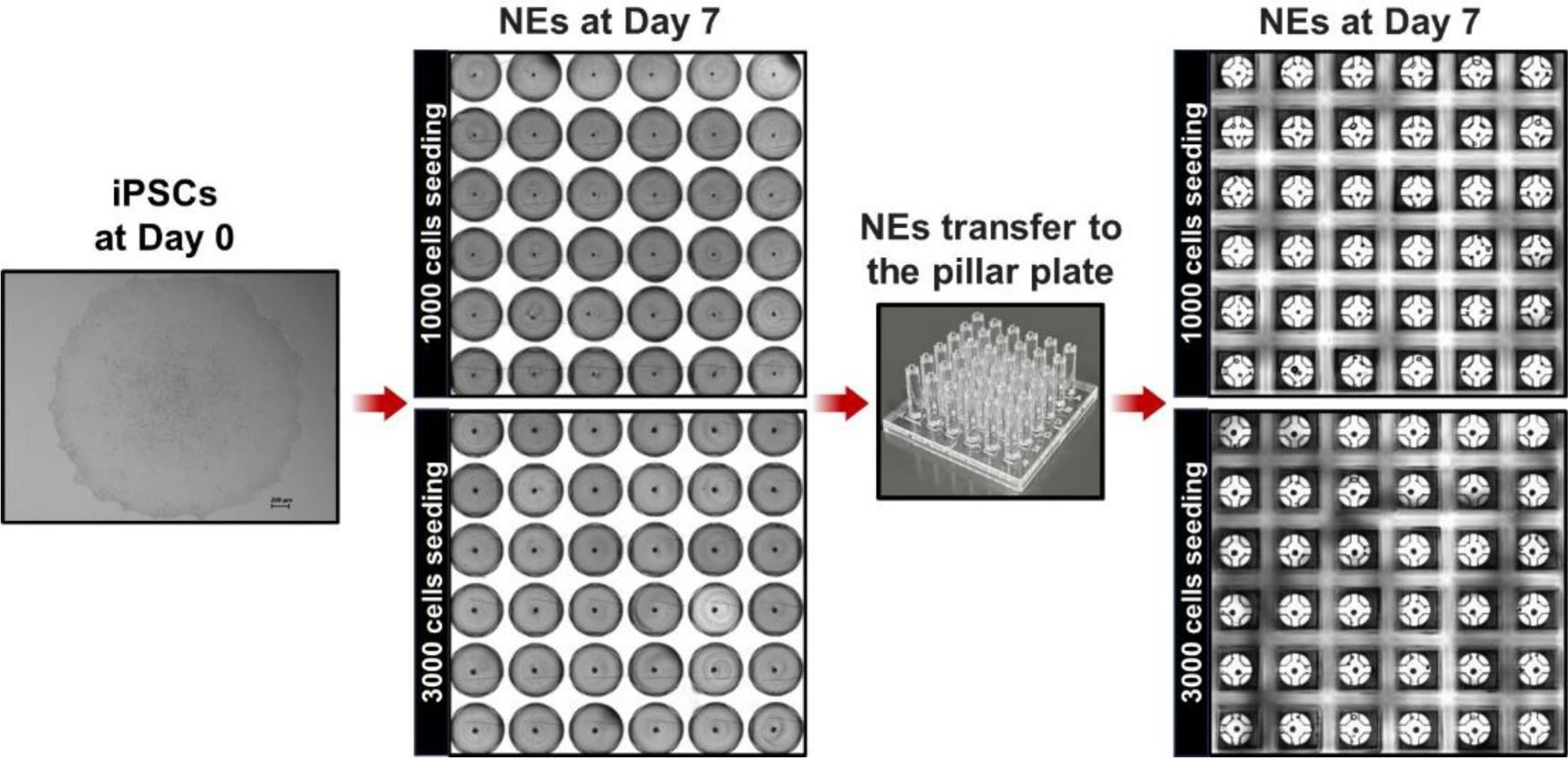
Neuroectoderms (NEs) formed in an ultralow attachment (ULA) 384-well plate at the seeding densities of 1,000 and 3,000 cells/well were transferred to a 36PillarPlate with Matrigel on day 7.

**Figure 2.**
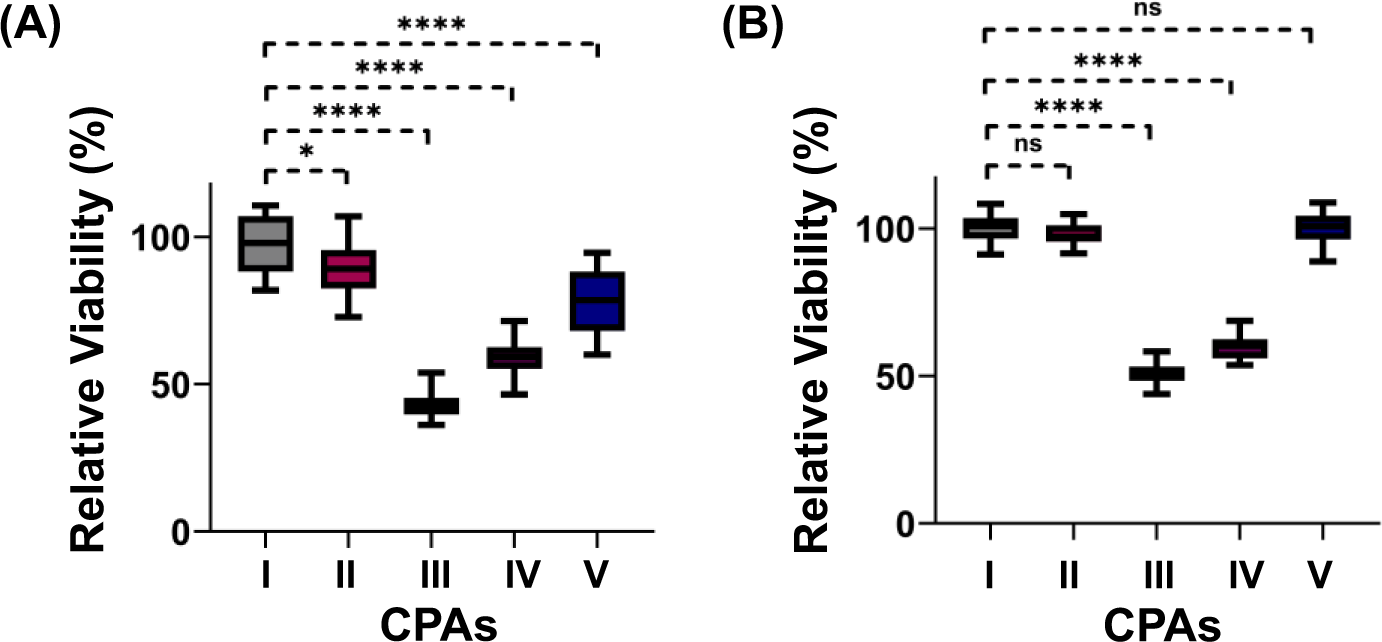
Relative viability of NEs after 2 hours of incubation at room temperature with cryoprotective agents (CPAs) including **(I)** Control (no CPA exposure), **(II)** PSC kit, **(III)** CryoStor, **(IV)** 3dGRO, and (**V)** 10% DMSO: **(A)** 1,000 and **(B)** 3,000 cell seeding density/well. The basal cytotoxicity of CPAs was compared with the control. The PSC kit and 10% DMSO maintained relatively high cell viability after 2 hours of incubation with the CPAs. n > 15. Statistical difference was analyzed with the control using one-way ANOVA. **** for p < 0.0001, *** for p < 0.001, ** for p < 0.01, * for p < 0.05, and ns = non-significant (p > 0.05).

### Estimating molecule diffusion into the core of NEs

Preventing ice crystal formation in the cores of spheroids and organoids with CPAs is crucial for enhancing cell viability post-cryopreservation. However, estimating the rate of CPA diffusion into the core of spheroids/organoids is challenging. To indirectly estimate this rate into the NE core, NEs on the pillar plate were incubated with CPA and calcein AM in a 384DeepWellPlate at room temperature for up to 2 hours, and changes in green fluorescence were monitored over time (**Figure 3**). Assuming similar diffusion rates for CPA and calcein AM, due to their small molecular sizes, it was necessary to incubate NEs with CPAs for 1 hour to achieve diffusion into the core. Calcein AM staining, which indicates cell membrane integrity, revealed that 3dGRO significantly reduces cell viability, while CryoStor causes NE size reduction. In contrast, neither the PSC cryopreservation kit nor 10% DMSO altered NE morphology or reduced cell viability.

**Figure 3.**
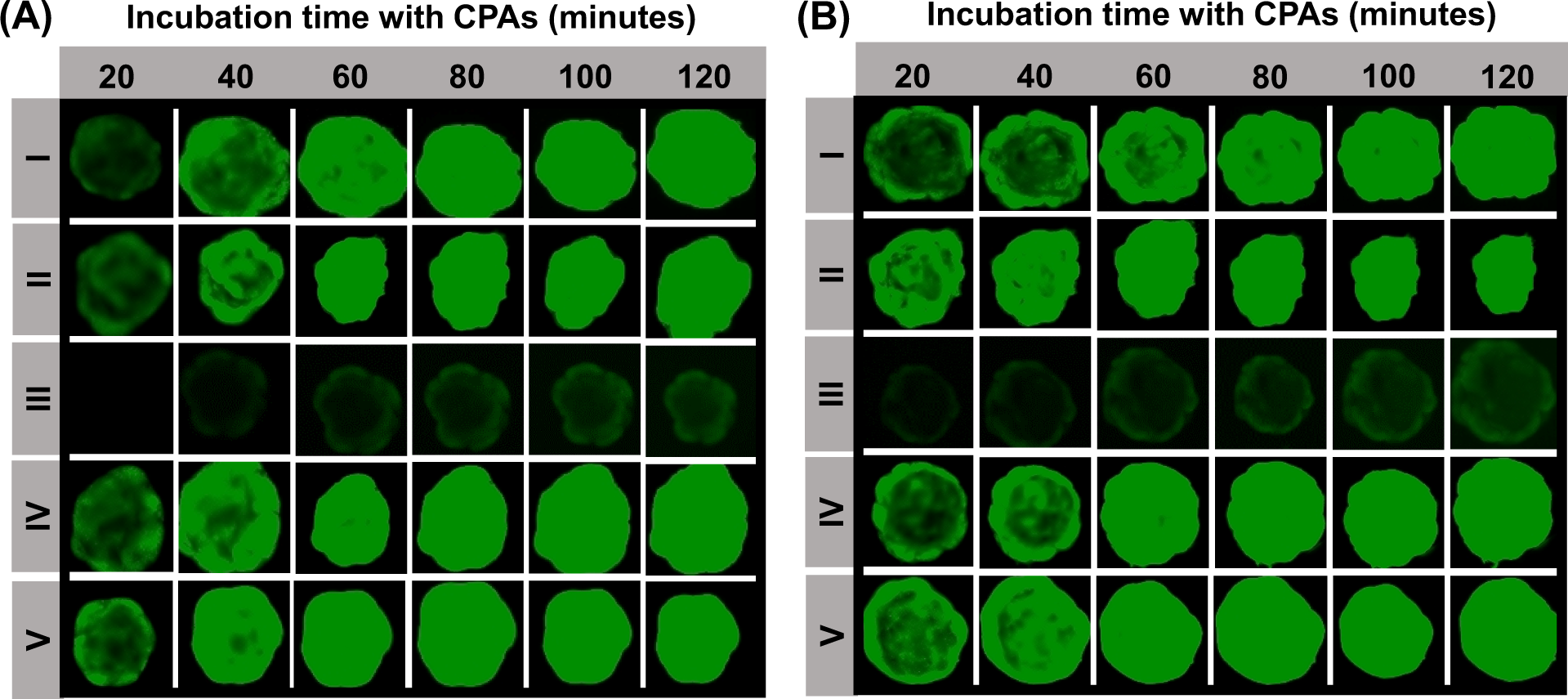
The staining of NEs with calcein AM in the presence of various CPAs, including **(I)** PSC kit, **(II)** CryoStor, **(III)** 3dGRO, **(IV)** 10% DMSO, and **(V)** Control (no CPA exposure). This was conducted for 2 hours at room temperature to evaluate the rate of molecule diffusion into the core of NEs and to assess the basal toxicity of the CPAs. The cell seeding density per well was **(A)** 1,000 and **(B)** 3,000. The diffusion of calcein AM into the core of NEs took approximately 1 hour, suggesting that the diffusion of CPAs may require at least the same amount of time.

### Changes in NE viability after incubation with the PSC cryopreservation kit

Cell viability may be altered after 24 hours post-exposure to toxic chemicals. Therefore, we assessed NE viability following exposure to the PSC cryopreservation kit for up to 2 hours, with subsequent incubation for 24 hours in a 5% CO_2_ incubator at 37°C (**Figure 4**). It was observed that NE viability decreased incrementally as the exposure time to the PSC cryopreservation kit increased. Nonetheless, NE viability was maintained above 80% for up to 80 minutes of exposure at room temperature. Consequently, the PSC cryopreservation kit was chosen for the cryopreservation of NEs on the pillar plate at - 80°C.

**Figure 4.**
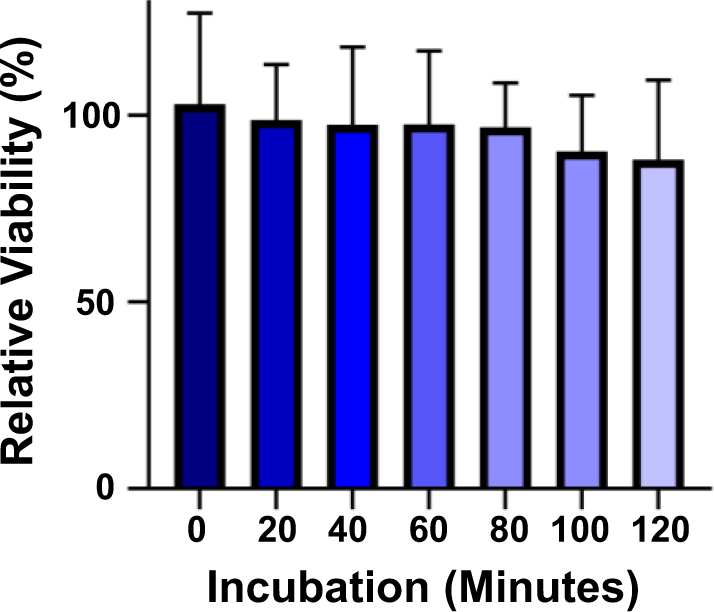
The changes in relative viability of NEs after a 2-hour incubation with the PSC kit at room temperature. Cell viability was subsequently assessed using the CellTiter-Glo luminescent cell viability assay kit following a 24-hour incubation period in a CO_2_ incubator. The number of replicates (n) was greater than 8 (n > 8).

### Determining optimal cell seeding density for NE cryopreservation

The diffusion of CPA into the core of NEs may be affected by the NE size, prompting an investigation into the influence of cell seeding density on cell viability. In brief, NEs created with initial cell densities of 1,000 and 3,000 cells on the 36PillarPlate underwent exposure to the PSC cryopreservation kit and were cryopreserved at - 80°C. Subsequently, a one-day incubation in NIM supplemented with RevitaCell was conducted, followed by the assessment of cell viability using the CellTiter-Glo 3D cell viability assay kit (**Figure 5**). As expected, the viability of NEs after cryopreservation with the PSC kit diminished as NE size increased, dropping from 58% to 36% in comparison to non-frozen NEs. These findings indicate that a lower cell seeding density, or a smaller NE size, is advantageous for preserving higher cell viability post-cryopreservation. Consequently, a cell seeding density of 1,000 cells has been selected for the continued optimization of the cryopreservation protocol.

**Figure 5.**
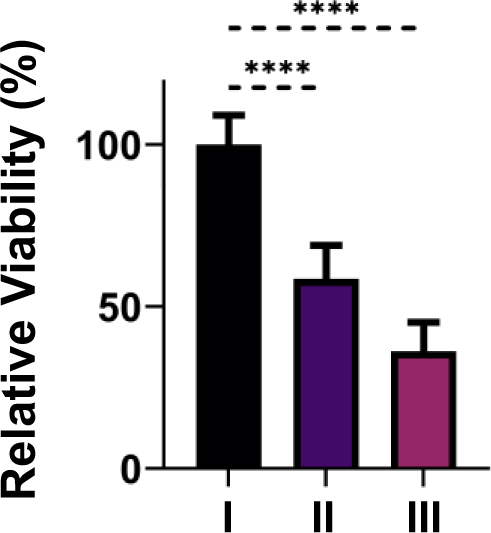
The changes in relative viability of NEs following cryopreservation using the PSC kit. The conditions compared were: **(I)** NE control (without cryopreservation) and NE samples after cryopreservation with the PSC kit at the seeding densities of **(II)** 1,000 cells and **(III)** 3,000 cells per well. The NEs were initially preserved at - 80°C overnight, followed by storage in liquid nitrogen for 3 days. Subsequently, they were thawed and incubated at 37°C for 24 hours. Post-incubation, cell viability was assessed using the CellTiter-Glo luminescent cell viability assay kit. n > 6.

### Cryopreservation of early-stage human brain organoids (ES-HBOs) on the pillar plate

After optimization of rinsing and recovery steps, we finalized the cryopreservation protocol for NEs on the pillar plate (**Figure 6**). The viability of the thawed NEs on the 36PillarPlate was measured by calcein AM and ethidium homodimer 1 (EthD-1) staining, which indicated highly viable NEs after 1 and 8 days post-cryopreservation and thawing with no apparent dead cells (**Figure 7A**). The morphology of the cryopreserved and thawed NEs on the 36PillarPlate was monitored for up to 30 days (**Figure 7B**). Expansion of the neuroepithelium was observed 4 days post-thawing, and the continued healthy morphology of the organoids was evident throughout the 30-day period.

**Figure 6.**
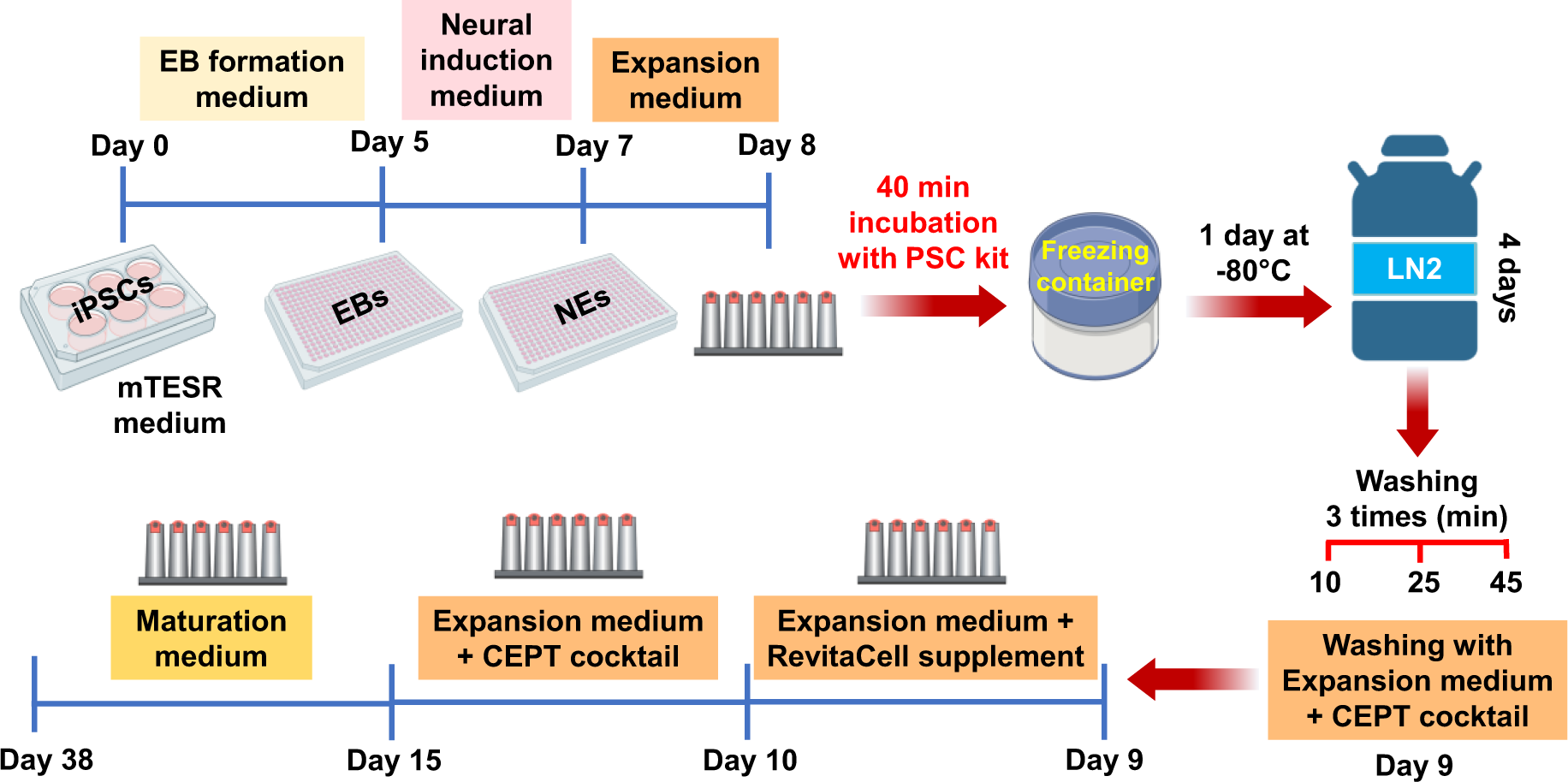
Optimum protocol of cryopreservation and thawing of NEs on the pillar plate.

**Figure 7.**
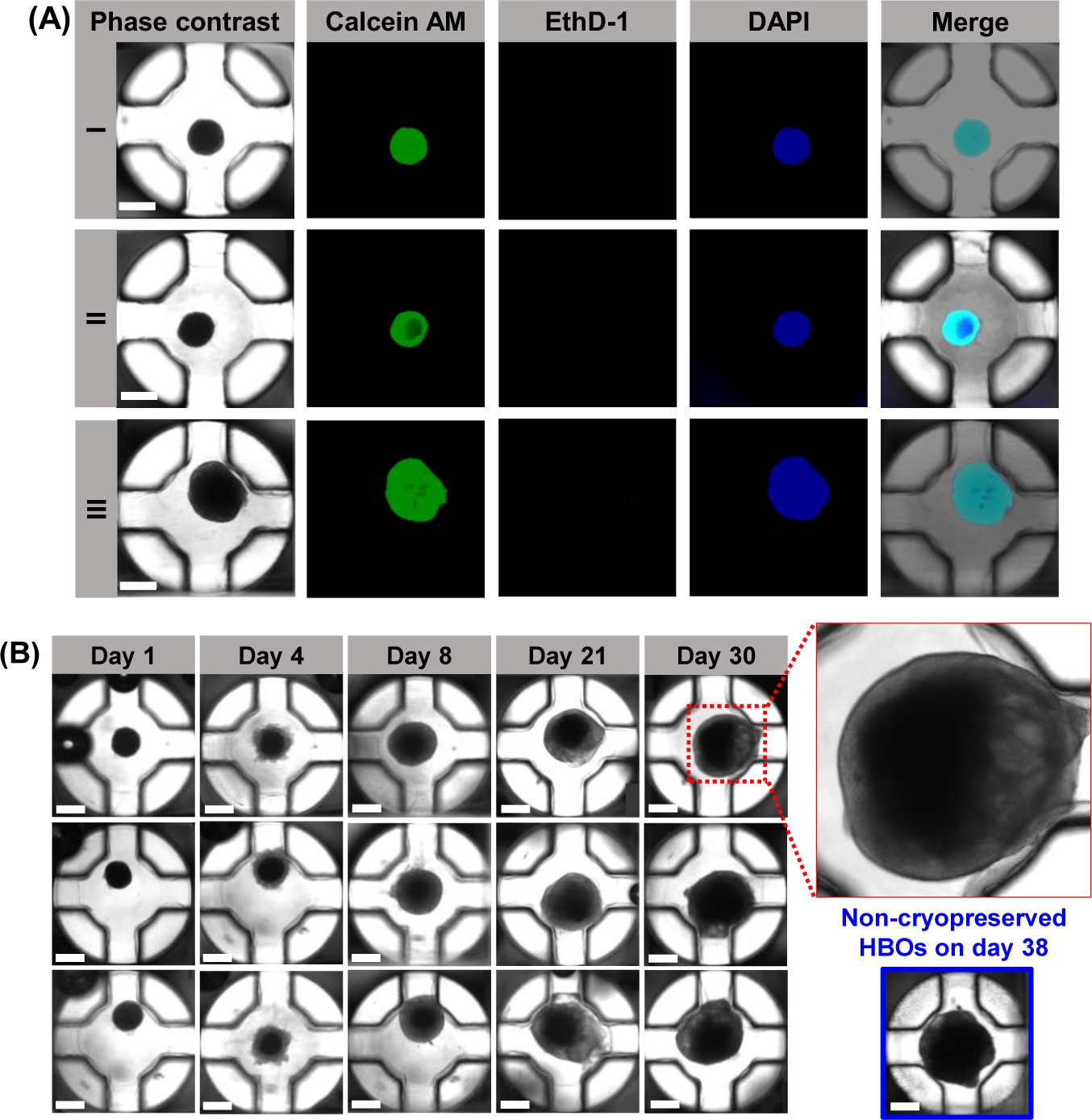
**(A)** The viability of NEs on the pillar plate before cryopreservation and after thawing: **(I)** Before cryopreservation, **(II)** 1 Day after thawing, and **(III)** 8 Days after thawing. The viability of the NEs was measured with calcein AM and ethidium homodimer 1 (EthD-1). Scale bars: 500 µm. **(B)** The changes in the morphology of NEs by differentiation on the pillar plate for 1, 4, 8, 21, and 30 days after cryopreservation and thawing. The cryopreserved NEs were properly recovered and showed the expansion of neuroepithelium on day 4 after differentiation. Scale bars: 500 µm.

Gene expression analysis by RT-qPCR indicated that the expression levels of the *OCT4* pluripotency marker, *PAX6* forebrain neuroprogenitor marker, and *FOXG1* forebrain marker in brain organoids derived from cryopreserved NEs were comparable to those in non-cryopreserved counterparts (**Figure 8**). On the other hand, the expression levels of the *SOX2* proliferating neuroprogenitor marker, *TBR2* intermediate progenitor marker, *TUBB3* neuronal cytoplasm marker, *CTIP2* deep cortical neuronal marker, and *MAP2* mature neuronal marker were slightly lower in brain organoids derived from cryopreserved NEs. These differences could potentially be ascribed to the recovery time necessary for the organoids after thawing, as well as the potential impact of cryopreservation on cell growth and differentiation. Furthermore, immunofluorescence staining of day 30 brain organoids from cryopreserved NEs showed high expression levels of SOX2, PAX6, TBR2, and MAP2 (**Figure 9**). In summary, cryopreserved NEs on the pillar plate successfully differentiated into brain organoids, maintaining gene/protein expression patterns similar to those of non-cryopreserved brain organoids.

**Figure 8.**
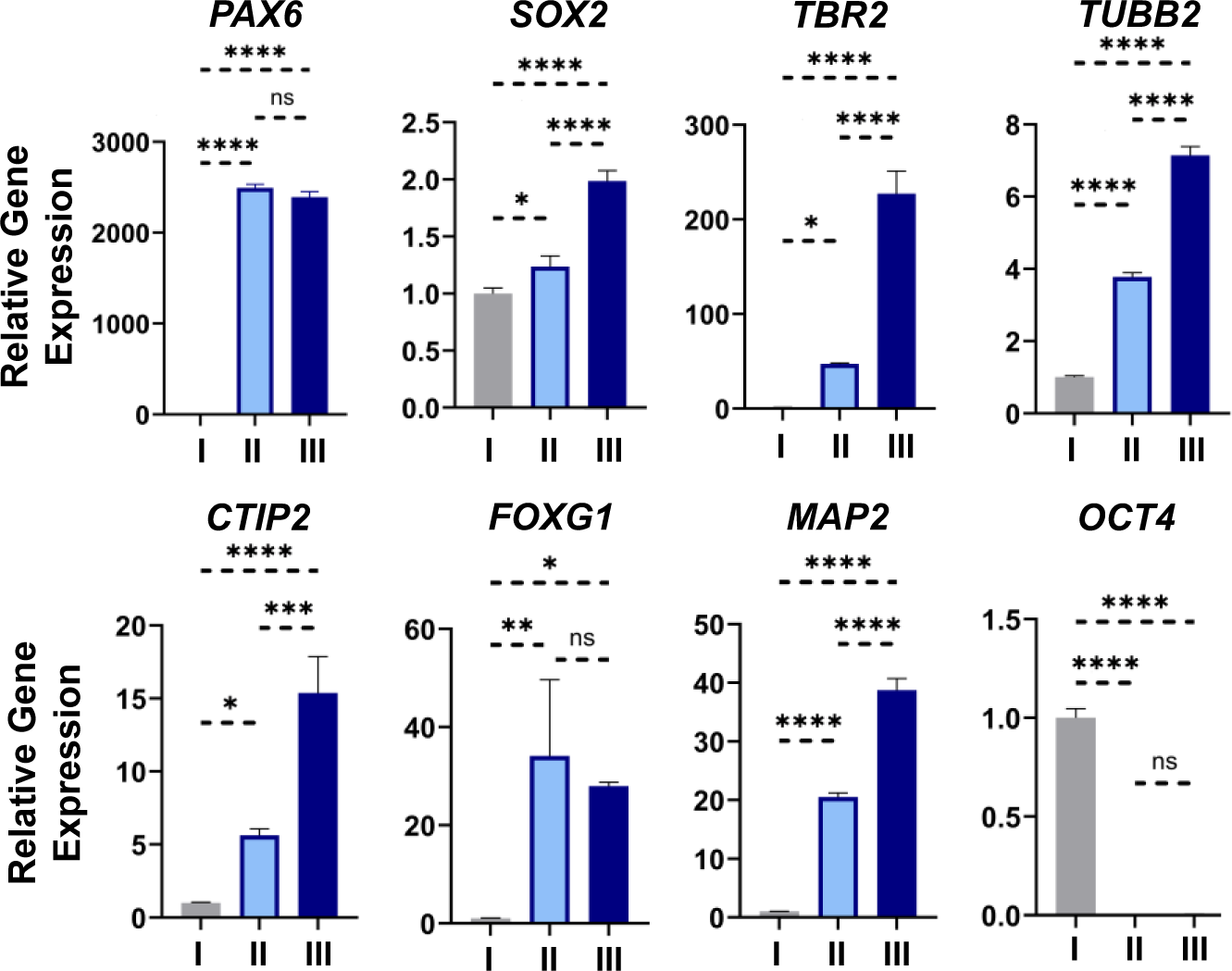
Characterization of gene expression in **(I)** iPSCs (control) and brain organoids differentiated from **(II)** cryopreserved and **(III)** non-cryopreserved NEs. Gene expression of *PAX6* forebrain neuroprogenitor marker, *SOX2* proliferating neuroprogenitor marker, *TBR2* intermediate progenitor marker, *TUBB3* neuronal cytoplasm marker, *CTIP2* deep cortical neuronal marker, *FOXG1* forebrain marker, *MAP2* mature neuronal marker, and *OCT4* pluripotency marker were analyzed by qPCR. Statistical significance was performed using one-way ANOVA. **** for p < 0.0001, *** for p < 0.001, ** for p < 0.01, * for p < 0.05, and ns = non-significance (p > 0.05). n = 10-12 per qPCR run.

**Figure 9.**
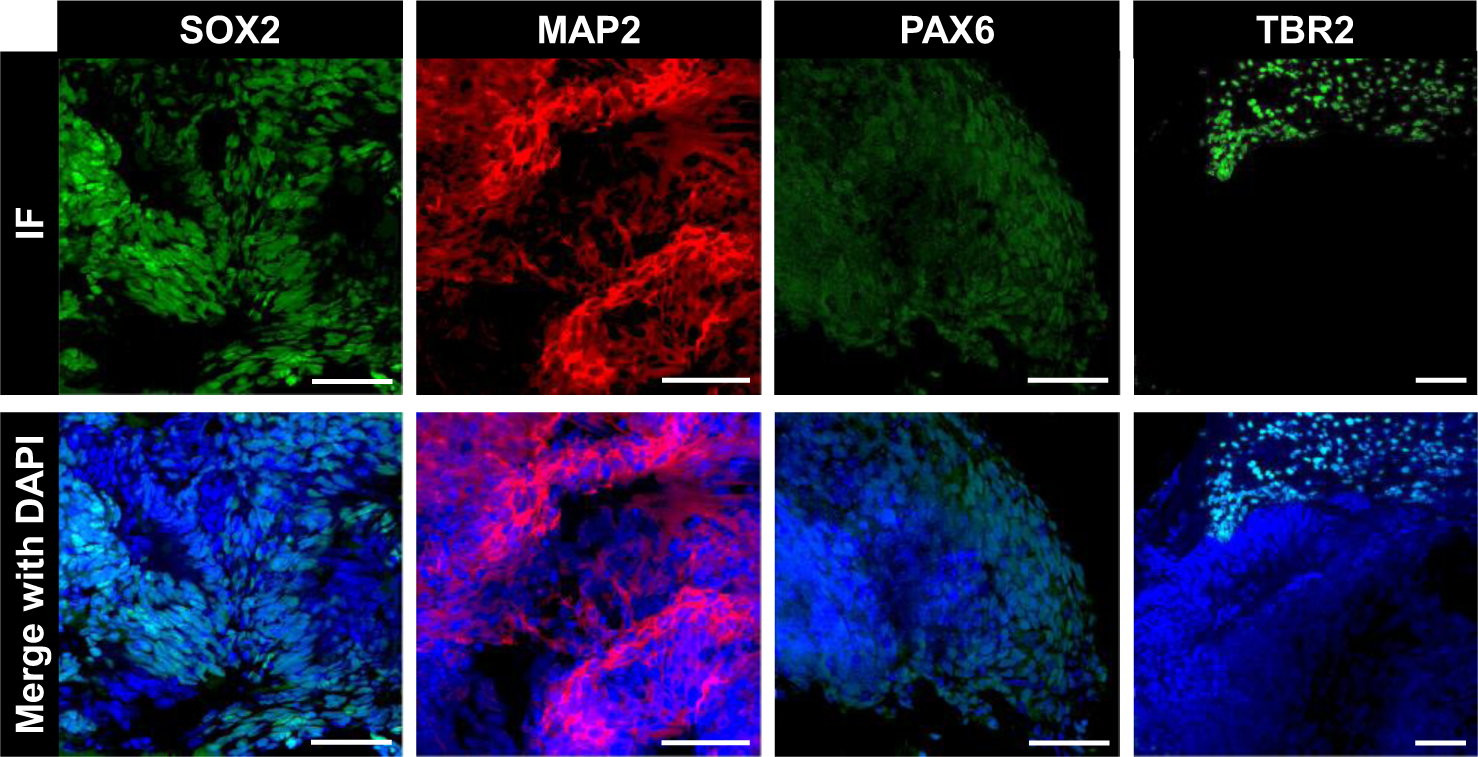
Immunofluorescence (IF) staining of brain organoids 30 days after thawing. SOX2 proliferating neuroprogenitor marker, MAP2 mature neuronal marker, PAX6 forebrain neuroprogenitor marker, and TBR2 intermediate progenitor marker were characterized in day 30 brain organoids differentiated from cryopreserved and thawed NEs. Scale bars: 50 µm.

## Discussion

Organoids have the potential to bridge the gap between 2D cell models and animal models, and they have been utilized for disease modeling, drug screening, and genetic engineering ^19,20^. The establishment of cryopreservation and thawing protocols for organoids could enhance biobanking and the utility of organoids in research and therapy. While several studies have investigated cryopreservation methods for 3D cell culture models, no specific protocol has yet been developed for brain organoids ^21–25^. The cryopreservation of organoids is influenced by various factors, including mechanical stress, cooling and thawing rates, organoid size, and the type of cryoprotective agents (CPAs). These factors significantly affect the viability of organoids after cryopreservation ^25,26^.

Traditional cryopreservation methods involve multiple steps, including organoid dissociation, centrifugation, and pipetting of the dissociated organoids ^27^, which could be reasons for cell loss and mechanical stress on cells. In contrast, *in situ* cryopreservation of organoids refers to the process of freezing organoids directly within their native environment, without removing them from the embedding matrix. This approach prevents cell loss and reduces mechanical stress compared to traditional cryopreservation methods. Liu *et al.* cryopreserved lung cancer organoids (LCOs) *in situ* in a microwell array. They demonstrated that *in situ* technology makes the freeze-thaw process more accessible, and LCOs maintained viability and structural integrity for drug sensitivity tests^28^. Additionally, embedding matrices such as Matrigel can protect cryopreserved cells and enhance cell viability during cryopreservation ^29^.

In this study, we developed optimal conditions for *in situ* cryopreservation of neuroectoderm (NE) in Matrigel and differentiation into brain organoids on the pillar plate. We adhered to the basic principle for successful cryopreservation, which is slow freezing and rapid thawing ^30,31^. During slow freezing, the penetration of cryoprotective agents (CPAs) into the core of 3D cells can be limited due to the typically low concentration of CPAs used. Therefore, we increased the incubation time with CPAs, enhancing their penetration into the core of NEs. However, CPAs can be toxic to cells, and prolonged exposure may impact cell viability. We assessed cell viability using calcein AM at various time points to evaluate the potential cytotoxicity of CPAs and their diffusion into the core of NEs. Of the four CPAs tested, CryoStor, 3dGRO, and 10% DMSO have been previously used for the cryopreservation of organoids, including human and animal intestinal, hepatic, and colon organoids ^32–34^. The efficacy of the PSC cryopreservation kit for brain organoid cryopreservation was examined for the first time in this work. The PSC cryopreservation kit demonstrated superior cell viability and organoid morphology among the four CPAs, leading to the successful cryopreservation of NEs on the pillar plate. Moreover, cryopreserved NEs on the pillar plate, when treated with a small amount of the PSC cryopreservation kit in the 384DeepWellPlate, resulted in rapid thawing of NEs and efficient CPA removal. Molecule diffusion and waste removal become less efficient as organoids increase in size, a common issue for the cryopreservation of organoids with a dense structure, such as brain organoids ^35,36^. Consequently, larger organoids pose more challenges for cryopreservation compared to smaller ones. In our work, NEs generated with a seeding density of 1,000 cells were chosen due to their relatively small size and adequate cell-cell interactions for brain organoid formation.

Cryopreserved NEs on the pillar plate maintained healthy morphology and differentiated into mature cerebral organoids. Nonetheless, a few days of cell recovery were necessary post-cryopreservation and thawing, allowing NEs to recover normal cell functions from the stress of freezing and thawing. Gene expression analysis and immunofluorescence staining confirmed that brain organoids derived from cryopreserved NEs were highly functional and could be used for organoid-based assays. For example, cryopreserved NEs on the pillar plate can be used to assess the impact of developmental neurotoxic compounds on normal brain organoid development.

## Conclusions

We have tested four commercially available CPAs, including the PSC cryopreservation kit, CryoStor, 3dGRO, and 10% DMSO, for the cryopreservation of NEs on the pillar plate. These NEs were successfully differentiated into mature brain organoids *in situ*. Among the CPAs tested, the PSC cryopreservation kit proved to be the most efficient, yielding high cell viability post-cryopreservation. The design of the pillar plate facilitated the immersion of small spheroids and organoids in CPA solutions, allowing for their cryopreservation in an empty 384-well plate. This method proved highly efficient for rapid thawing and CPA removal, crucial for maintaining high cell viability. We anticipate that the pillar plate could be utilized for the *in situ* cryopreservation of various spheroids and organoids, potentially streamlining the cryopreservation process and subsequent assays, thereby enhancing the throughput and reproducibility of organoid-based assays.

## Acknowledgment

This study was supported by the National Institutes of Health (NCATS R44TR003491, NIDDK UH3DK119982, and NIEHS R43ES035653).

## Supplementary Information

**Supplementary Table 1.**
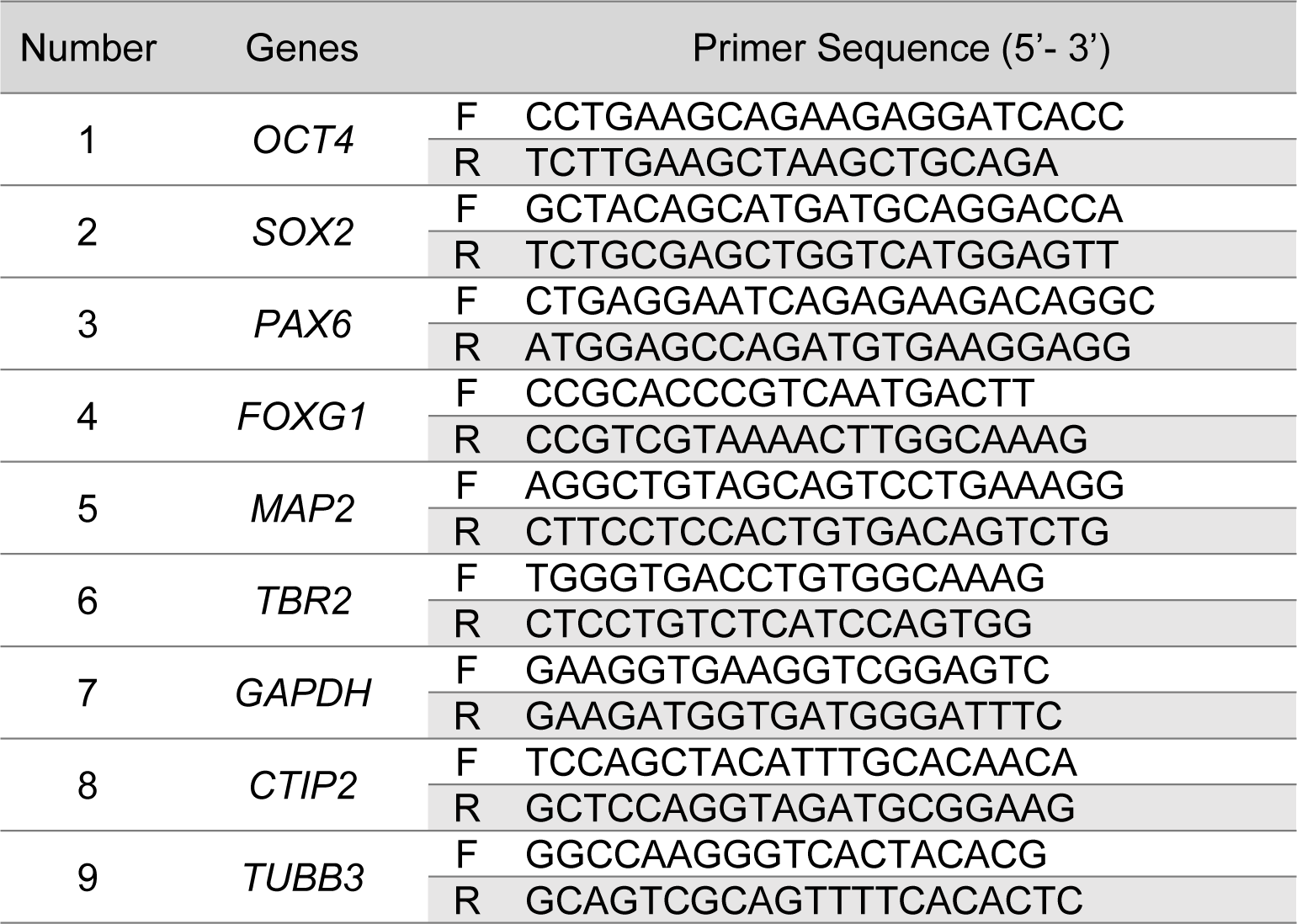
List of primers.

**Supplementary Table 2.** List of primary antibodies.

| Name | Species | Dilution | Vendor | Catalog # |
| --- | --- | --- | --- | --- |
| PAX6 | Rabbit | 1:350 | Abcam | ab195045 |
| SOX2 | Mouse | 1:40 | Santa Cruz Biotech | sc-365823 |
| TBR2 | Rabbit | 1:200 | Abcam | 216870 |
| MAP2 | Rabbit | 1:200 | Abcam | 183830 |

**Supplementary Table 3.** List of secondary antibodies.

| Name | Dilution | Vendor | Catalog # |
| --- | --- | --- | --- |
| Donkey anti-Rabbit IgG (H+L) Highly Cross-Adsorbed Secondary Antibody, Alexa Fluor 647 | 1:200 | ThermoFisher | A-31573 |
| Donkey anti-Mouse IgG (H+L) Highly Cross-Adsorbed Secondary Antibody, Alexa Fluor 488 | 1:200 | ThermoFisher | A-21202 |
| Donkey anti-Rabbit IgG (H+L) Highly Cross-Adsorbed Secondary Antibody, Alexa Fluor 488 | 1:200 | ThermoFisher | A21206 |

